# Inhibition of Microbial Beta-Glucuronidase Does Not Prevent Breast Carcinogenesis in the Polyoma Middle T Mouse

**DOI:** 10.1101/746602

**Authors:** Amanda P. Beck, Hao Li, Samantha M. Ervin, Matthew R. Redinbo, Sridhar Mani

## Abstract

**Purpose:** To demonstrate whether inhibition of intestinal microbial beta (β)-glucuronidase (BGUS) abrogates tumor formation in a MMTV-PyMT mouse breast cancer model.

**Methods:** Female MMTV-PyMT heterozygote mice (4 weeks old) were randomized to oral gavage with vehicle or UNC10201652 (20 μg/day), a microbial BGUS inhibitor, for 9 weeks. The entire animal carcass was assessed for tumor deposits by histology and immunohistochemical staining for tumor (Ki67, PCNA) and breast specific (ER, PR, Cyclin D1, aSMA, Integrin β1) markers.

**Results:** The MMTV-PyMT breast pathology in our study simulates prior published reports of tumor incidence and aggressiveness. There was no significant difference in the tumor histology, number of tumors (lesions), and patterns of spread of tumors in the UNC10201652 versus the vehicle treated mice. Similarly, there were no significant differences in the semi-quantitative scores for expression of ER, PR, Ki67, PCNA, or Integrin β1. There were also no major differences seen in qualitative screening of Cyclin D1 and aSMA.

**Conclusions:** MMTV-PyMT mice administered UNC10201652, when compared to vehicle treated mice, show a similar abundance of breast tumor (and tumor initiating) lesions ranging from hyperplasia to frank carcinoma. There is a trend, however small, that the incidence of hyperplastic and adenomas may be decreased in UNC10201652 treated mice. Further refined dosing strategies in MMTV-PyMT are planned to clarify its biological significance. To our knowledge this is the first report of use of any BGUS inhibitor in chemoprevention of breast tumors using a genetic model simulating human breast cancer.

## Introduction

Breast cancer afflicts one in eight women and is second to lung cancer in women with respect to cancer deaths [1]. Hormone receptor-positive breast cancer constitutes the vast majority of cases of breast cancer world-wide [2, 3]. Recently, it has been proposed that the human intestinal tract houses bacteria that may intrinsically convert metabolized estrogen (in the form of estrogen glucuronide) to the parent hormone, estrogen, via actions of microbial specific enzymes, beta (β)-glucuronidase (BGUS) [4, 5]. This axis of re-purposing parent estrogen in normal physiology, termed the “estrabolome”, is a likely mechanism to keep estrogen levels fine-tuned in our bodies [4, 6]. The evidence supporting this concept is demonstrated by human studies whereby the intestinal microbiome composition affects the levels of systemic estrogen[7–10]. Fecal BGUS activity in women correlate with reduced levels of plasma concentration of estrogen [11]. In a recent paper, using conjugated estrogens and bazedoxifene, manipulation of fecal BGUS activity was achieved to lower systemic estrogen levels in mice to levels commensurate with what might be alleviate menopause-associated symptoms in women [10]. In addition, nipple aspirate microbial BGUS activity (as well as associated microbial species) were significantly elevated in breast cancer cases versus healthy controls [12]. Studies of chemically-induced mammary carcinogenesis in mice suggest that diets that lower microbial BGUS activity in cecal contents are associated with lower incidence of mammary tumor, albeit with no change in circulating estrogen levels [13]. Indeed, microbial BGUS enzymes have been implicated in carcinogen metabolism, which could confound interpretation of these models in terms of BGUS activity and protection against breast carcinogenesis [14, 15]. Overall, there is compelling human data suggesting a link between microbial BGUS activity either in feces and/or nipple aspirates with systemic estrogen availability and cancer association.

This link has not been investigated systematically in non-chemically induced (i.e. genetic) models simulating human breast carcinogenesis. Specifically, no genetic models of breast cancer in rodents have been analyzed with intestinal BGUS modulating compounds. In this regard, a well-studied mouse model of breast tumor progression, the MMTV-PyMT^634^ mice, is endowed with the development of early lesions (hyperplasia/adenoma) that are strongly estrogen-receptor positive, and this was a key driver of our present study [16, 17]. This mouse model has been extensively and reproducibly characterized with respect to dietary interventions and tumor progression stages and hormone-receptor status and other biomarkers [16, 18–25]. With respect to our hypothesis, if the intestinal “estrabolome” had significant estrogen metabolic effects, it might be conceivable that the development of early lesions (e.g., hyperplasia/adenoma) might drive lesions forward towards a frank carcinoma due to the availability of recirculating estrogen. However, specific strategies would then be needed to ablate intestinal BGUS in order to demonstrate a cause-and-effect of the “estrabolome” hypothesis.

In contrast with clinical and rodent “association” studies that have been published to-date, it has not been feasible to prove “causation” of effect when microbial-specific BGUS is ablated *in vivo*. In this regard, until recently, there were no specific inhibitors or strategies available to reduce microbial BGUS in any organ and thus show its direct effect on reducing breast carcinogenesis. Recently, though, BGUS inhibitors have been developed that are potent, selective toward microbial GUS enzymes, and non-lethal to microbial cells [26]. These inhibitors were initially employed to block the reactivation of the active metabolite of the solid tumor chemotherapeutic irinotecan in the intestinal lumen, and were shown to significantly reduce gut damage and resultant diarrhea in mice [27, 28]. The active metabolite of irinotecan, SN-38, is glucuronidated to inactive SN-38-G in the liver and other protective tissues by Phase II drug metabolizing UGT enzymes [29]. SN-38-G is sent to the gut for excretion, where it is subject to reactivation by microbial BGUS, leading to sufficient SN-38 in the GI tract to cause intestinal damage and the dose-limiting side effect of irinotecan, delayed diarrhea. The initial compound characterized, Inhibitor 1, was shown to have no effect on circulating plasma levels of irinotecan or its key metabolites [30]. Inhibitor 1 was also shown to block small intestinal ulcers caused by the non-steroidal anti-inflammatory drugs diclofenac, indomethacin and ketoprofen, which are all glucuronidated and subject to reactivation by microbial BGUS [31–34]. Inhibitor 1 was found to have no effect on plasma levels of diclofenac in mice [31]. Finally, in 2018, it was shown that Inhibitor 1 blocks anastomotic leakage after intestinal resection in rats given diclofenac [35]. Thus, a potent and specific BGUS inhibitor that does not kill microbial cells has been shown to be effective in blocking several negative outcomes with drugs or surgical interventions.

An improved BGUS inhibitor, UNC10201652, was recently described that highjacks the catalytic cycle of the BGUS active site, utilizing a piperizine warhead to form a long-lived covalent inhibitor-glucuronide bound to the active site of the enzyme [36]. As such, this compound is more potent than Inhibitor 1, as demonstrated recently in detailed analyses of several human gut microbial BGUS enzyme orthologs, their processing of drug and non-drug substrates, and the inhibition of these reactions [37, 38]. Thus, we chose to examine the effects that UNC10201652 has on the development of breast tumors in the PyMT mouse model of hormone receptor positive tumorigenesis.

## Methods

### Mice

MMTV-PyMT^634^ heterozygote male mice were a generous gift from Jeffrey Pollard PhD (Albert Einstein College of Medicine, Bronx, NY) [17] and bred (crossed to female FVB, code 27, Charles River) in our vivarium. At each successive breeding generation, all pups (male and female) were genotyped and separated in cages (after weaning) based on gender. Female (4 weeks old) mice generated from each breeding pair were randomly allocated to one of two treatment groups (n = 15/treatment group; total n = 30 female mice generated over a 24 month period) using a coin-toss approach as previously utilized [39]. Cage allocation was also determined by coin toss and to maintain homogeneity in the microbiome exposure, within each treatment group, mice were randomly shuffled between their respective treatment group cages prior to treatment group assignment (e.g., mouse 1 on day 1 was in cage 1, then on day 2 allocated to cage 3 within its respective treatment group etc.). Mice were treated in metabolic cages to limit communal coprophagy. The control group consisted of oral gavage with vehicle (0.67 % DMSO/0.9% saline; 100 μl) and the treatment group, UNC10201652 (Inh9, 20 μg/day) gavaged (in 0.67 %DMSO/0.9% saline; 100 μl), each starting day 1 week 3 after weaning (4 week old mice) and consecutively dosed daily for 9 weeks. Dosing was omitted over weekends (2 days) but resumed each starting weekday (Monday). Gavage was necessary as prior studies in our laboratory showed that mice are averse to consuming UNC10201652in water due to poor taste. UNC10201652 is also poorly soluble in water. Mice under 4 weeks of age cannot tolerate gavage with UNC10201652.

Mice were fed standard laboratory chow (Lab Diet, #5058) and animal facility water as provided for all barrier facility mice. They followed a standard 12h sleep-wake cycle and all cages were housed in the same room and in the same rack. Fecal pellets were collected from each mouse at baseline and at necropsy. Euthanasia and distress relief intervention was conducted strictly according to the approved Institute for Animal & Use Care Committee of the Albert Einstein College of Medicine, Bronx, NY (IACUC# 20161019) and the Army Medical Research and Materiel Command (USAMRMC) Animal Care and Use Review Office (ACURO) (#BC161093) protocols. Mouse breeding and genotyping was conducted under a separate protocol approved by the Institute for Animal & Use Care Committee of the Albert Einstein College of Medicine, Bronx, NY (IACUC#: 20160701; renewed breeding IACUC#00001036).

### Genotyping

Mice were genotyped exactly as previously described and published and using primer sets as described in above [40]. The representative PCR gel for mouse genotyping and classification is shown in Supplementary Figure 1. The genotyping primer sets for PCR:

Forward 5’ – GGAAGCAAGTACTTCACAAGGG – 3’ and

Reverse 5 ‘ - GGAAAGTCACTAGGAGCAGGG – 3’

The expected band size is 556 bp (Supplementary Figure 1).

### Histopathology

For each mouse, all mammary glands were collected. Two of the mammary glands, as well as all lung lobes, were assigned for histopathologic examination by a board-certified veterinary pathologist (APB). Tissues for histopathology were fixed for 48 hours in 10% neutral buffered formalin and then routinely processed for paraffin embedding, sectioning, and staining by the Albert Einstein Cancer Center Histology and Comparative Pathology Facility. Each sample was sectioned at three levels (separated by 100 μm) in duplicate (i.e. each level had 2 slides) for assessment of lesion morphology. The lesions determined included hyperplasia, adenoma, early carcinoma and late carcinoma. Histopathological examination of the mammary tumors of each mouse was conducted and when present, lesions were categorized as hyperplasia, adenoma/mammary intraepithelial neoplasia (MIN), early carcinoma and late carcinoma as per previous work with PyMT mice [17].

Slides were H&E stained using a standard protocol and imaged with a Zeiss AxioImager A.2 microscope and Zeiss Axiocam 305 color using ZEN software. The histologic lesions satisfying the definitions of hyperplasia, adenoma, early and late carcinoma were then individually ascertained across an entire slide and their areas (defined by a percentage of the grids covered) were expressed as an approximate percentage (%) of the total area of the mammary gland on that slide.

### Immunohistochemistry

Immunohistochemical staining was performed using the following antibodies: estrogen receptor alpha (ERα) (1:1000; 106132-T08, Sino Biological), progesterone receptor (PR), Her2/Neu (1:250; C-18, Santa Cruz Biotechnology), cyclin D1 (1:250; 72-13G, Santa Cruz Biotechnology), α-smooth muscle actin (α-SMA)(1:200; D4K9N; Cell Signaling), integrin-β1 (1:50; 4B7R, Santa Cruz Biotechnology), PCNA (1:2000; PC10, Invitrogen), and Ki67 (1:400; D3B5, Cell Signaling). All immunohistochemistry was performed on formalin-fixed, paraffin-embedded mammary tissues, and appropriate positive and negative controls were used for each antibody. Antigen-retrieval methods were used for all antibodies. The detection method was either ImmPRESS HRP Anti-Rabbit IgG (MP-7451; Vector Laboratories) or ImmPRESS HRP Anti-Mouse IgG (MP-7452; Vector Laboratories), according to the species source of the antibody. The chromogen was DAB (SK-4100; Vector Laboratories) and the counterstain was Harris hematoxylin for all antibodies.

Assessment of staining for ERα, PR, Her2/Neu, cyclin D1, α-SMA, and integrin β1 was performed semi-quantitatively as follows: 0 = no staining; 1 = rare/minimal positive staining; 2 = occasional positive staining; 3 = moderate amount of positive staining; 4 = widespread positive staining. One slide (20 high-power fields) per mouse was assessed. For Ki67, the percentage of positively stained neoplastic cells was evaluated according to published recommendations [41], which suggest the assessment of at least three randomly selected high-power fields (HPF), providing that staining is homogenous across the section. The percentage of positively stained neoplastic cells for PCNA was also assessed across three randomly selected HPF (400x).

### Statistical Analysis

The published literature of chemoprevention studies using the MMTV-PyMT mouse model [19, 22, 42] suggests that potential drugs that could have chemoprevention properties have a *standardized* effect size (SES) that range between 20-80% (0.2 to 0.5). The assumption of our analysis is based on the fact that ~90-100% of the MMTV-PyMT mice at 12-14 weeks develop tumors covering the entire spectrum of histology evaluated [17]. If we assume that over a 24-month period, we can efficiently generate and study 30 mice (~ based on breeding we estimate obtaining 1-2 females per month and conducting the experiment in a staggered manner), to show an SES of ≥ 55% assuming a common standard deviation (30%) we would require 24 mice totally (distributed in a 1:1 treatment allocation manner). This assumes a two-tailed analysis of outcome with 95% power and *a* = 0.05 (https://www.stat.ubc.ca/~rollin/stats/ssize/n2.html). The experiment inflation factor (loss of mice prior to start of randomization like premature demise) is ≤20% (0.2), then the total sample size required would be 30 mice [43]. An intent-to-treat analysis was conducted on all mice and outcomes.

One-way analysis of variance (1-Way ANOVA) as well as Two-way analysis of variance (2-Way ANOVA), with Sidak’s test for multiple comparisons conducted in swapped fashion, as well as diagnostics for residuals using Spearman’s rank correlation test for heteroscedasticity and for gaussian distribution of residuals, were calculated using GraphPad Prism version 8.2.0 (272) for macOS (GraphPad Software, La Jolla, California, USA).

## Results

### The Morphology of MMTV-PyMT mouse tumors

All mice examined (in both control and treatment groups) exhibited similar histologic findings. Mammary tissue contained a range of abundant proliferative lesions, from hyperplasia to adenoma/MIN to carcinoma. Hyperplastic foci consisted of ducts lined by one to several layers of jumbled epithelial cells that were polygonal, with indistinct cell borders, a small amount of pale amphophilic cytoplasm and round vesicular nuclei with variably prominent nucleoli. Anisocytosis and anisokaryosis (pleomorphism) were minimal and mitoses were rare. The hyperplastic ducts were surrounded by a distinct and intact basement membrane. Foci of adenoma/MIN were characterized by a more florid epithelial proliferation than hyperplasia, but still confined by an intact basement membrane (no invasion). The acini and ducts were filled and expanded by epithelial cells with similar characteristics as in the hyperplastic foci, though the cells in adenoma /MIN exhibited slightly more anisocytosis, anisokaryosis and mitoses (up to 3/400x field). There were frequent coalescing foci of carcinoma, in which epithelial cells showed features as described above in adenoma/MIN but with increased pleomorphism and mitoses (6-10/400x field). Importantly, there was also a lack of distinct/intact basement membrane with invasion of the neoplastic cells into adjacent stroma/fibroadipose tissue. Multifocally, there was frequent and abundant central necrosis within the carcinomas. There were also foci that exhibited features of late stage carcinoma, with solid sheets of epithelial cells with no remaining acinar structures visible. The lung metastases were evaluated both grossly and by microscopy. They were qualitatively similar in the number of lesions, the area of lung involvement and size in both the vehicle and UNC10201652 treated groups (Figure 1 A-E).

**Figure 1.**
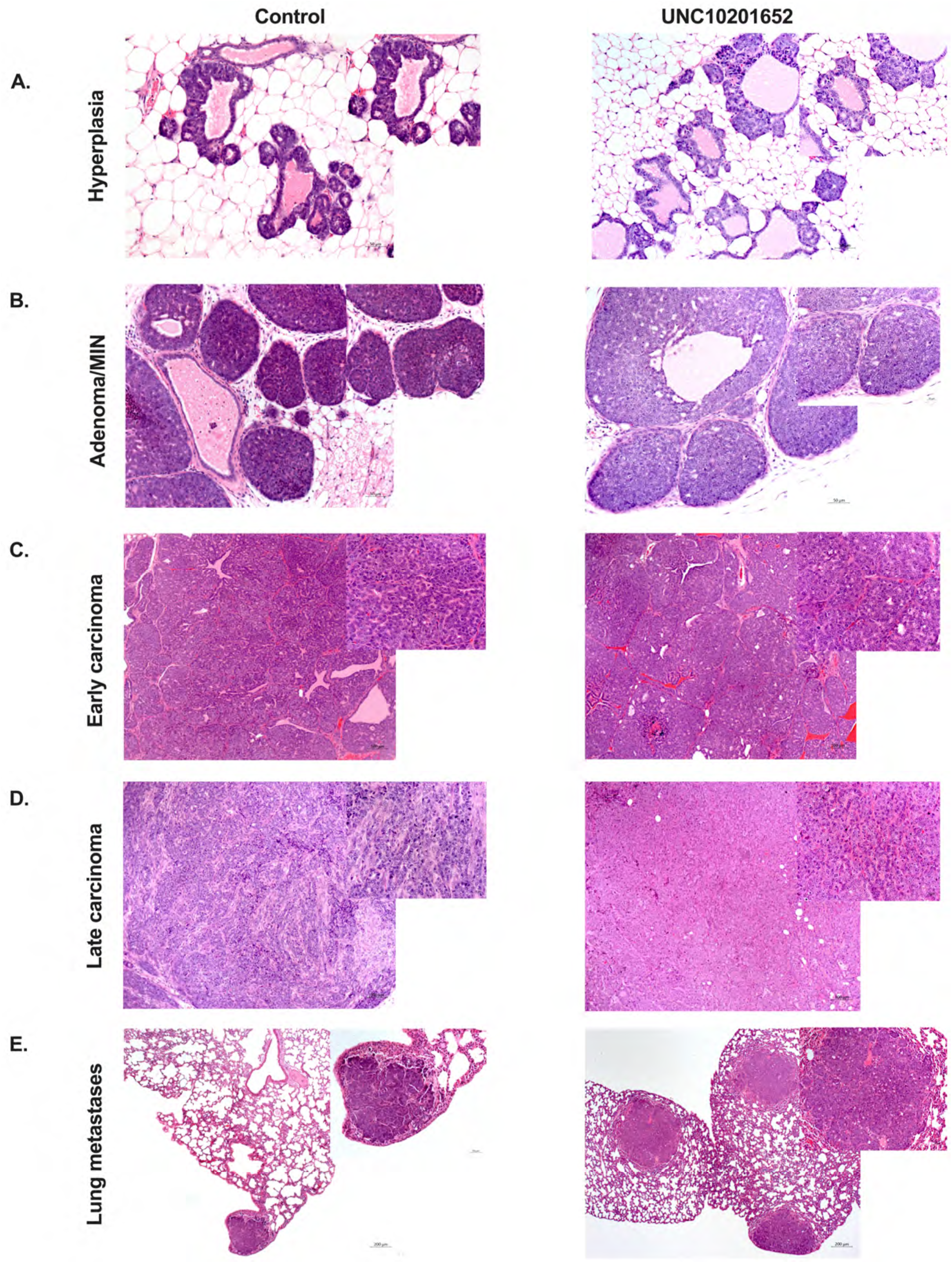
Histology of MMTV-PyMT tumors. Mammary tissue of all mice (vehicle and UNC10201652 treated) contained a range of abundant proliferative lesions, from hyperplasia to adenoma/MIN to early and late carcinoma. A) Hyperplastic foci consisted of ducts lined by one to several layers of jumbled epithelial cells with minimal pleomorphism. The hyperplastic ducts were surrounded by a distinct and intact basement membrane. Representative H&E stain, 200x. B) Within foci of adenoma/MIN, acini and ducts were filled and expanded by epithelial cells with slightly more anisocytosis, anisokaryosis and mitoses than in hyperplastic regions. They remained confined by an intact basement membrane. Representative H&E stain, 200x, inset (right upper panel): 400x. C) Early carcinomas exhibited neoplastic cells with increased pleomorphism and mitoses, as well as invasion of the neoplastic cells through the basement membrane and into adjacent stroma/fibroadipose tissue. Representative H&E stain, 100x, inset: 400x. D) Late stage carcinomas were composed of solid sheets of pleomorphic epithelial cells with no remaining acinar structures visible and a high mitotic index. H&E stain, 100x, inset: 400x. E) Metastatic foci within the lungs were composed of neoplastic cells with characteristics as described for the primary mammary neoplasms. They were qualitatively similar in the number of lesions, the area of lung involvement and size in both the vehicle and UNC10201652 treated groups. Representative H&E stain, 50x, inset: 200x.

### UNC10201652 does not alter the tumor incidence in MMTV-PyMT mice

The entire mouse carcass was dissected and *all* mammary glands removed. The assigned veterinary pathologist (APB) randomly picked two mammary glands and lungs for histologic reading at three levels cut into the prepared tissue block. The levels were numbered serially (1, 2 and 3) and two percentage counts (2 slides evaluated at each level) for each tumor histology (hyperplasia, adenoma, early carcinoma and late carcinoma) was generated. The minimum % area of hyperplasia, adenoma, early carcinoma and late carcinoma, were not significantly different between the control and UNC10201652 treated group, respectively (Figure 2A-C). Similarly, the maximum % area of hyperplasia, adenoma, early carcinoma and late carcinoma, were also not significantly different between the control and UNC10201652 treated group, respectively (Figure 2D-F).

**Figure 2.**
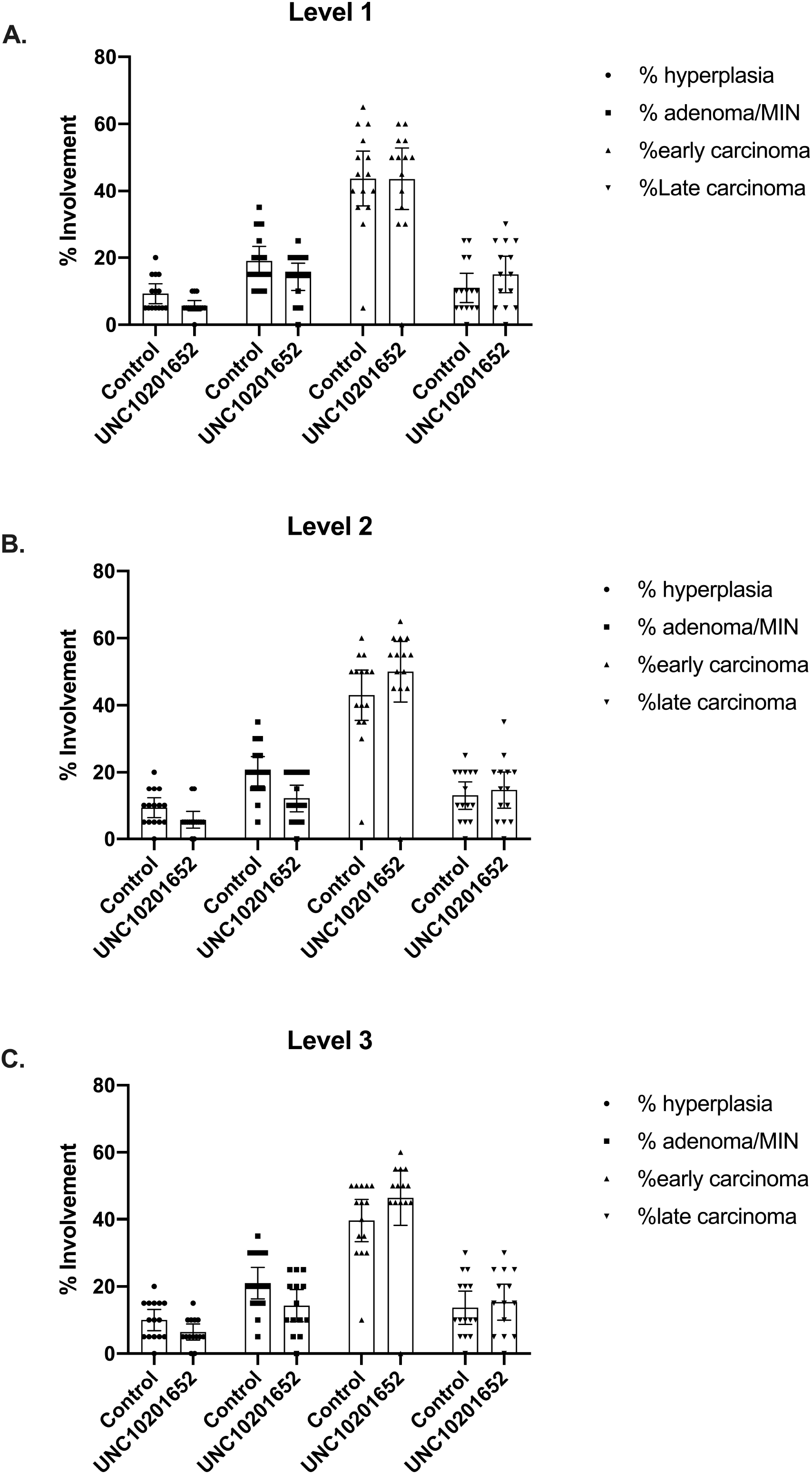
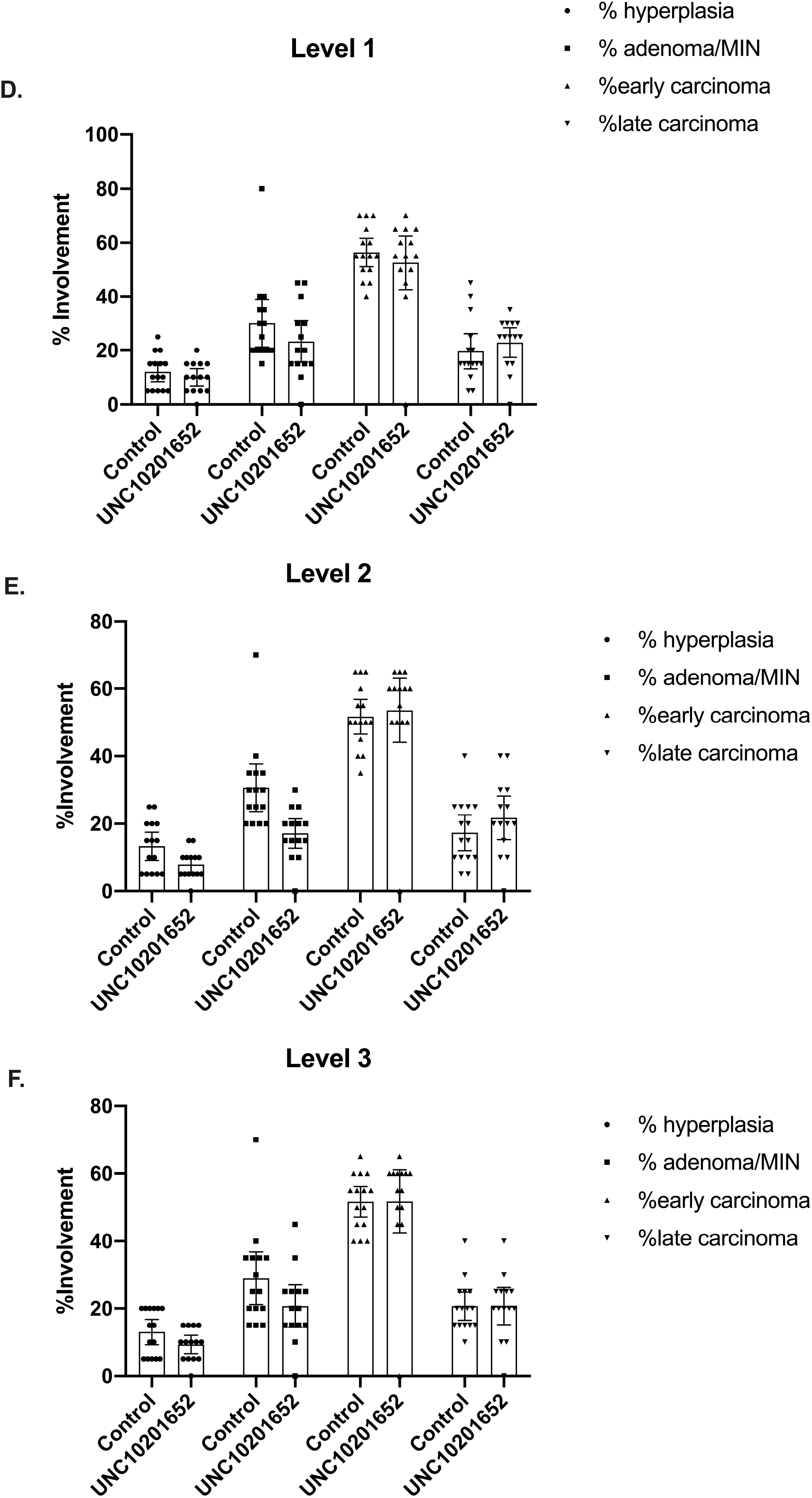
Effect of UNC10201652 on breast tumor incidence in MMTV-PyMT female mice. Female mice (4 weeks old) were administered either control vehicle or UNC10201652 by oral gavage daily for 9 weeks. Three levels of tissue were cut and 2 slides per level was evaluated for a minimum and maximum percent (%) area of involvement of any given lesion (hyperplasia, adenoma, early carcinoma, late carcinoma). A) Level 1 B) Level 2 C) Level 3, minimum % area involvement by lesion type (n = 15 mice/treatment group). D) Level 1 E) Level 2 F) Level 3, maximum % area involvement by lesion type (n = 15 mice/treatment group). There was no significant difference in any tumor type between treatment groups (Two-way ANOVA followed with Sidak’s test, p > 0.1) at any level evaluated.

### UNC10201652 does not alter the breast tumor marker distribution in MMTV-PyMT mice

#### Estrogen Receptor (ERα)

ER is expressed largely in hyperplastic and adenomatous lesions in MMTV-PyMT tumors [17]. The tumor and surrounding mammary tissues were assessed for ER expression in our mice. The semi-quantitative scoring of positive staining was analyzed (Figure 3A, upper panel) and represented numerically. The expression score is significantly lower in the carcinoma stage as compared with either hyperplasia or adenoma (p < 0.0001, Two-way ANOVA followed with Sidak’s test). There is no significant difference within each lesion type between the two treatment groups for the extent of ER-positivity (Figure 3A, lower panel).

**Figure 3.**
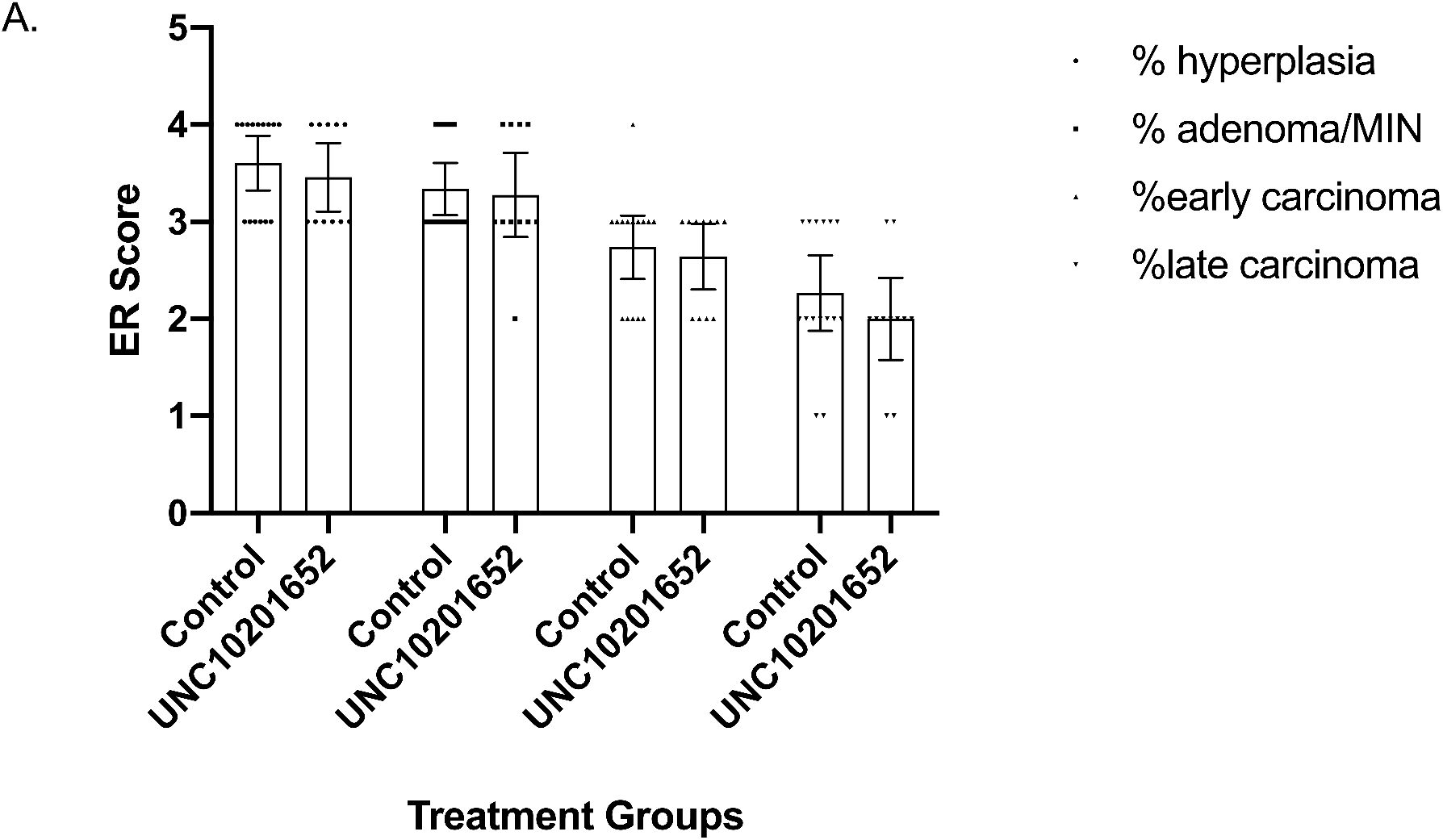
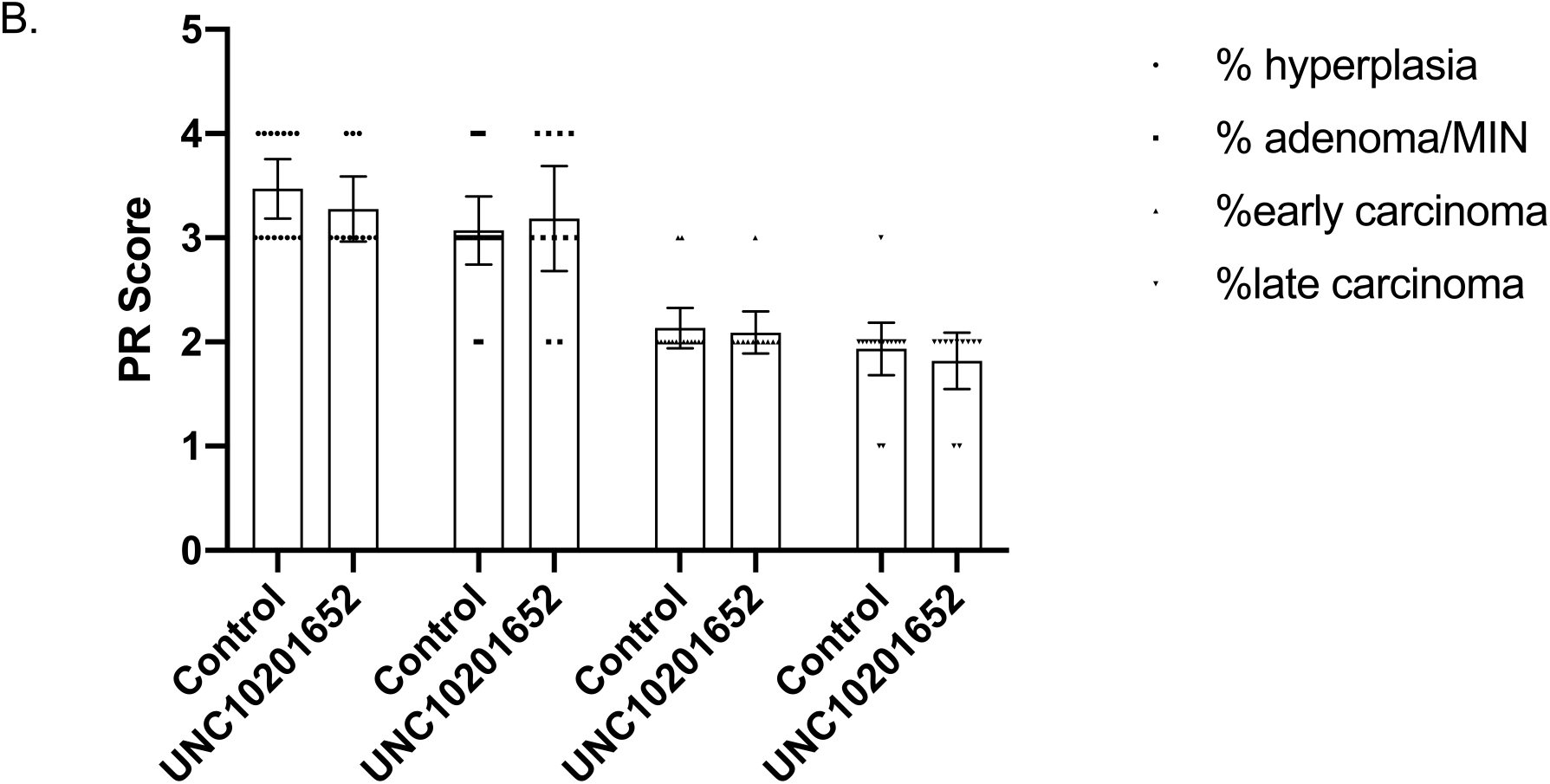
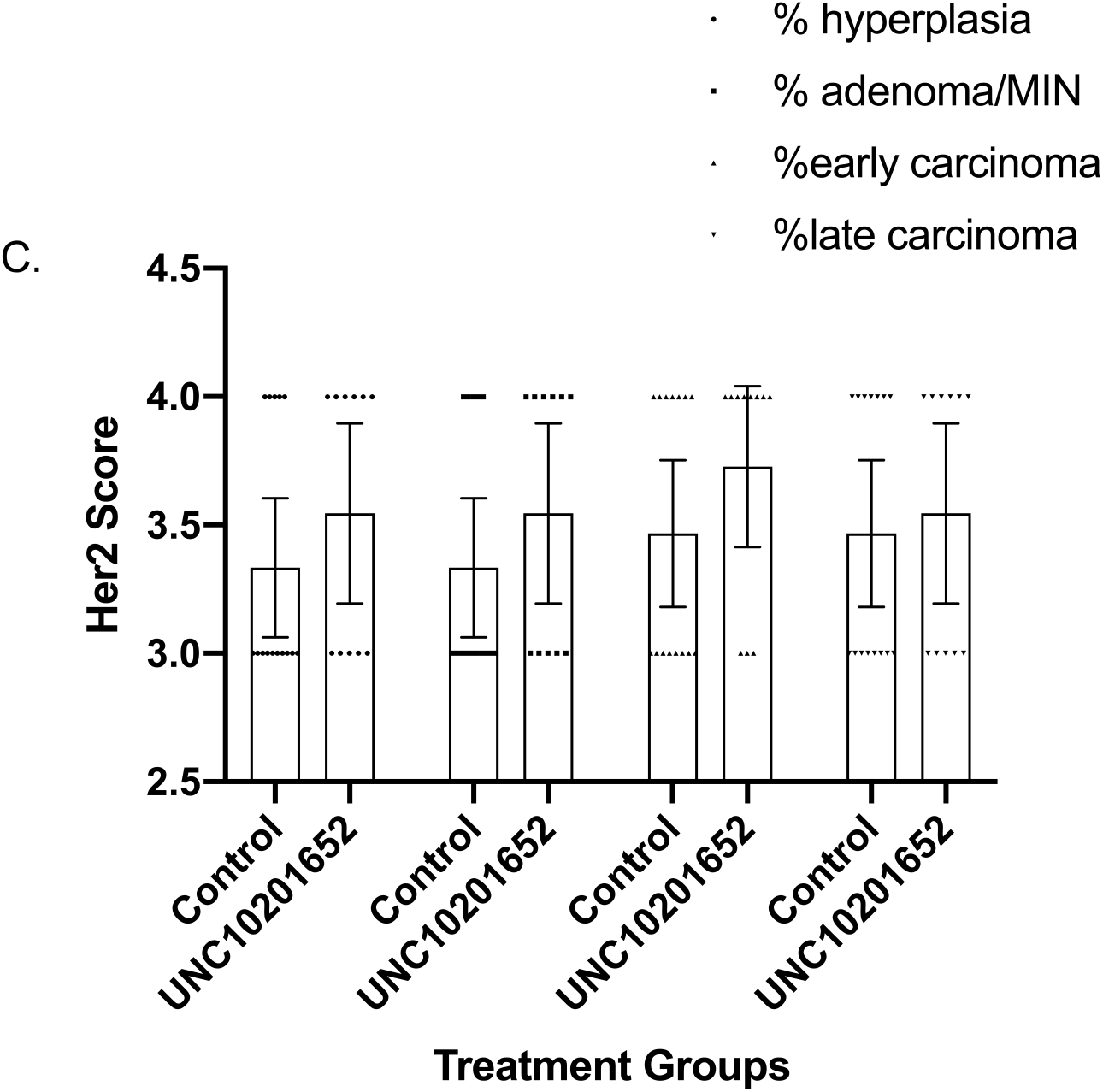
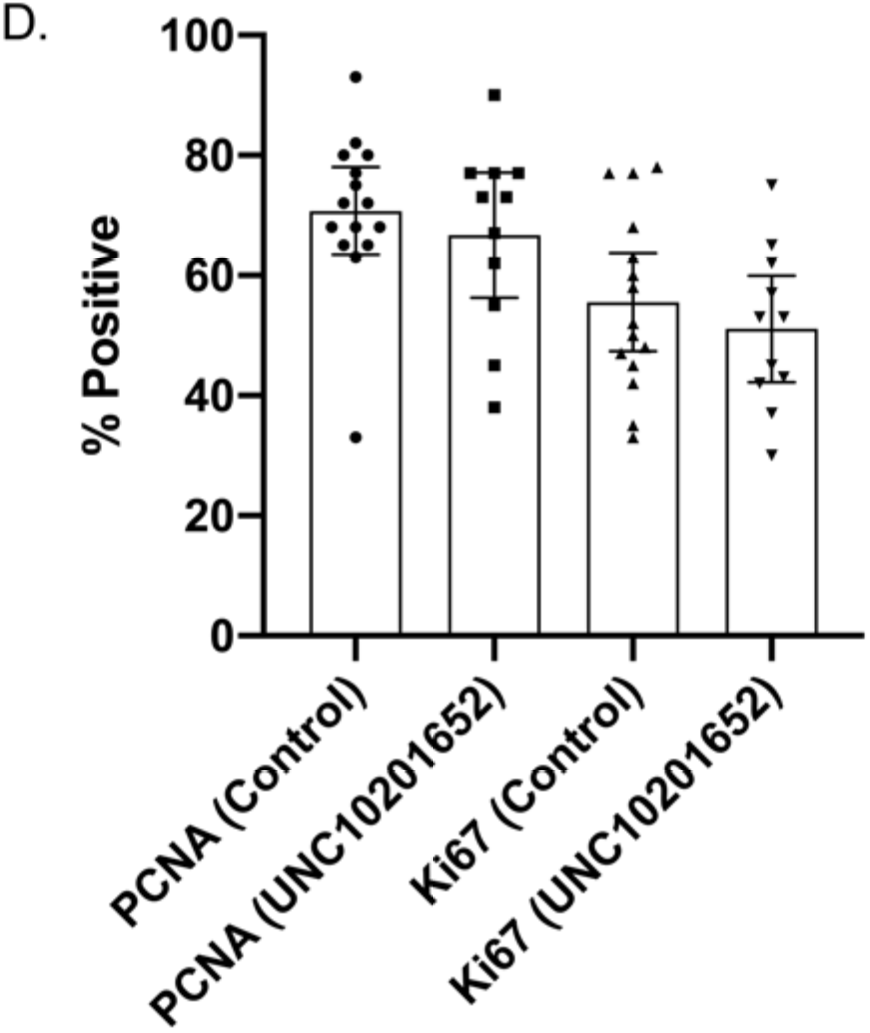
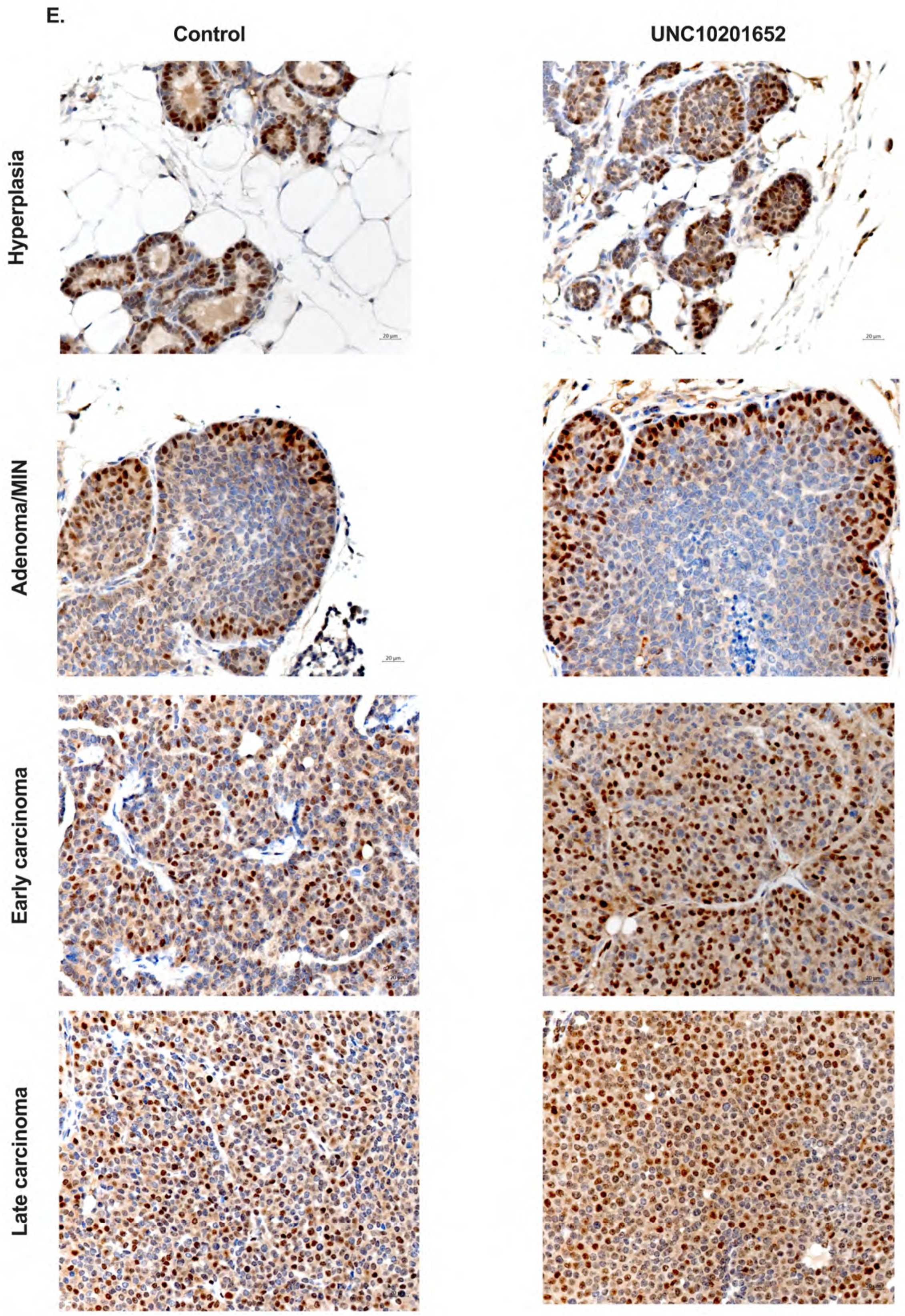
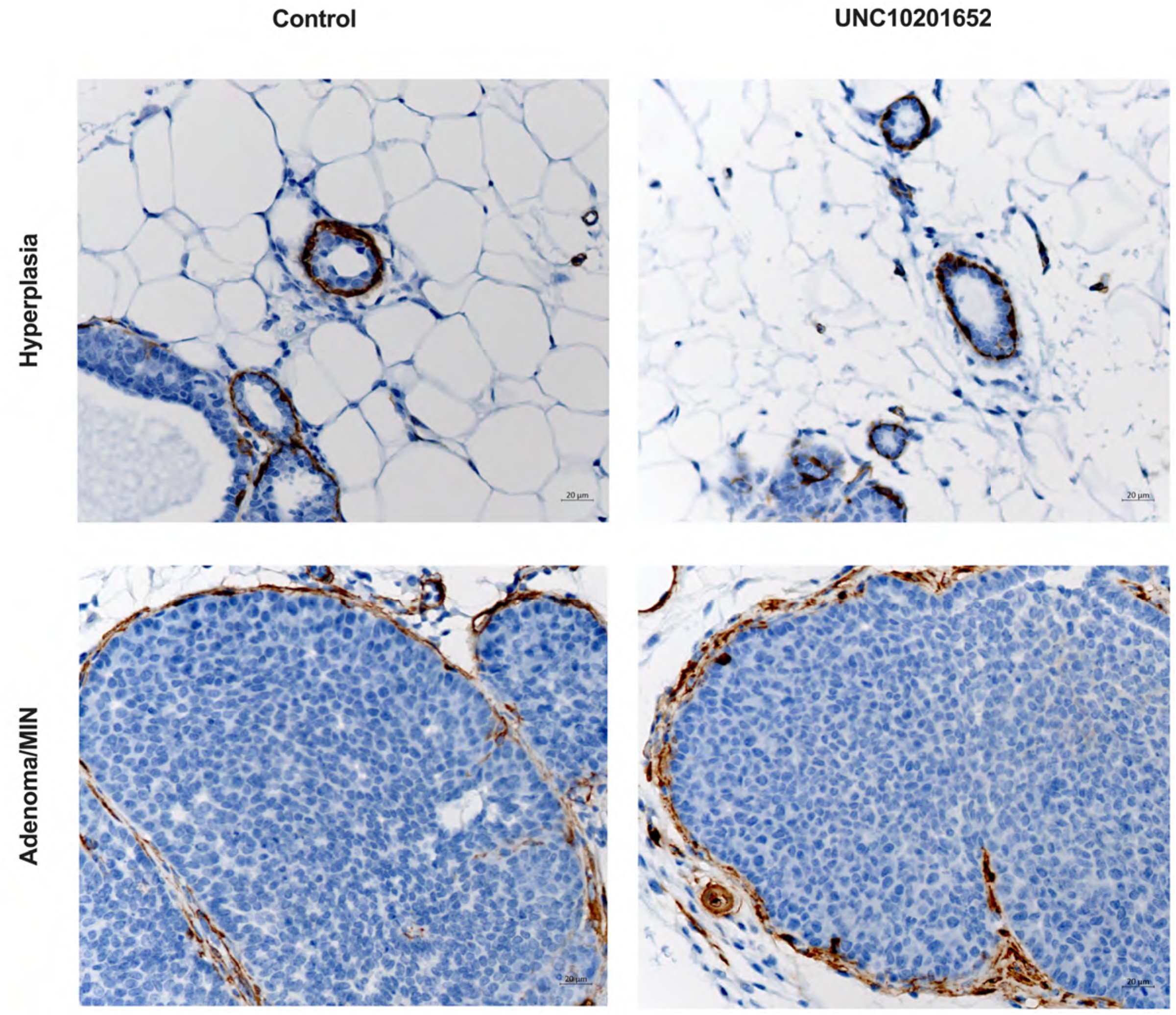
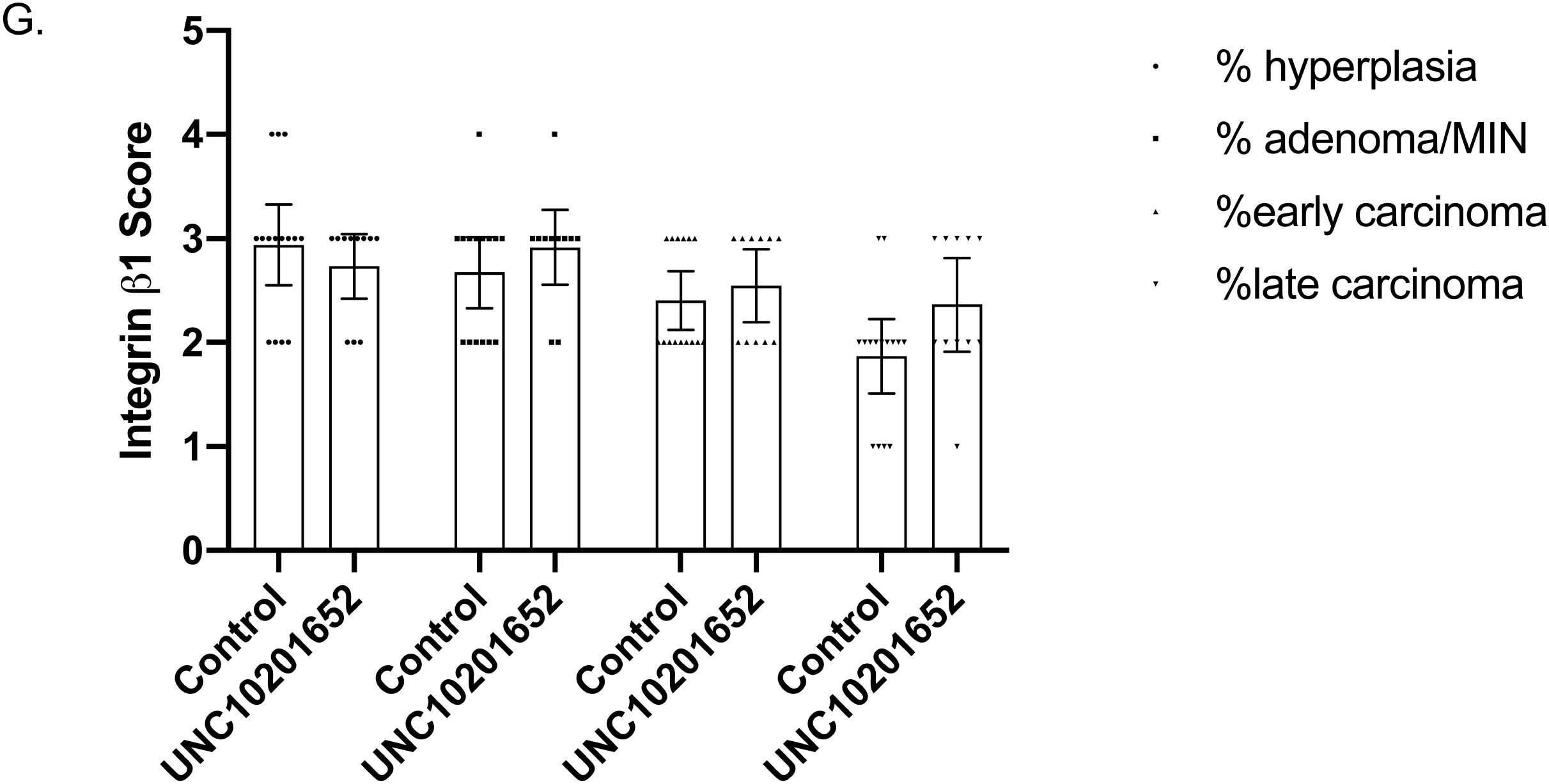
Immunohistochemical Staining of MMTV-PyMT tumors. Female mice (4 weeks old) were administered either control vehicle or UNC10201652 by oral gavage daily for 9 weeks. The semi-quantitative scores are plotted for control (n = 15 mice) and UNC10201652 (n = 11) for A) the estrogen receptor alpha (ERα)*, B) the progesterone receptor (PR)*, C) Her2/neu*, D) Ki67**, PCNA**, E) Cyclin D1 (400x), F) α-SMA (400x), and G) Integrin β1*. *There was no significant difference in any tumor type between treatment groups (Two-way ANOVA followed with Sidak’s test, p > 0.1) at any level evaluated. ** There was no significant difference in any tumor type between treatment groups (One-way ANOVA followed with Sidak’s test, p > 0.1) at any level evaluated.

#### Progesterone Receptor (PR)

Similar to ER, PR is expressed largely in hyperplastic and adenomatous lesions in MMTV-PyMT tumors [17]. Additional staining was conducted using antibodies targeting PR. The semi-quantitative scoring of positive staining was analyzed (Figure 3B, upper panel) and represented numerically. The expression score is significantly lower in the carcinoma stage as compared with either hyperplasia or adenoma (p < 0.0001, Two-way ANOVA followed with Sidak’s test). As with ER, there was no significant difference within each lesion type between the two treatment groups for the extent of PR-positivity (Figure 3B, lower panel).

#### Her2/neu

Her2/neu is expressed is low abundance in hyperplasia and its score intensity rises with the progression of the lesion reaching highest levels in late carcinomas [17]. For Her2/neu, the semi-quantitative scoring of positive staining was analyzed (Figure 3C, upper panel) and represented numerically. However, unlike prior published data [17], we were unable to find significant differences in the semi-quantitative scoring between lesion types in our mice p = 0.647, Two-way ANOVA followed with Sidak’s test). By visual inspection, there was a trend, however, for increased Her2/neu expression in the more advanced stages beyond adenomas within each treatment group. There was no significant difference within each lesion type between the two treatment groups for the extent of PR-positivity (Figure 3C, lower panel).

#### Ki67

Ki67 is a reliable marker for testing proliferation in early and late breast tumor lesions (~40-85% positive staining depending on the lesion type) in the MMTV-PyMT mouse (albeit C57BL/6J background) model [44]. In this analysis, we looked at all lesions in a given HPF in a slide and the semi-quantitative “average” scoring of positive staining (in 3 random HPF) was analyzed (Figure 3D, upper panel) and represented numerically. Our staining percentages ~ 35-77% are similar in quality and percent of cells expressing Ki67-positivity to published data [44]. There was no significant difference within each lesion type between the two treatment groups for the extent of PR-positivity (Figure 3D, lower panel).

#### Proliferating Cell Nuclear Antigen (PCNA)

Similar to Ki67, for PCNA staining a semi-quantitative “average” scoring of positive staining (in 3 random HPF) was analyzed (Figure 3D, upper panel) and represented numerically. There was no significant difference within each lesion type between the two treatment groups for the extent of PR-positivity (Figure 3D, lower panel).

#### Cyclin D1

Cyclin D1 is robustly elevated as lesions progress from hyperplasia to frank carcinoma and the distribution of the positive staining within the lesion varies [17]. At the hyperplasia and adenoma/MIN stage, cyclin D1 positively stained cells were mainly located at the outer margins of the proliferative lesions, while at both early and late carcinoma stages, there were increased numbers of cyclin D1-positive cells within the tumors, as well as distribution of the positive cells throughout the entire tumor (Figure 3E).

#### α-SMA

Immunostaining for α-SMA was utilized to stain myoepithelial cells. In hyperplastic lesions, similar to normal/unaffected mammary duct, myoepithelial cells form a single complete layer upon which luminal epithelial cells sit. Adenoma/MIN lesions were surrounded by a single layer of myoepithelial cells, though there were multiple foci in which the layer appeared incomplete. In more advanced lesions (carcinomas), the myoepithelial cell layer was completely disrupted and lost. (Figure 3F).

#### Integrin β1

For Integrin β1, which is only marginally expressed in MMTV-PyMT tumors and decreases as the lesion advances [17], the semi-quantitative scoring of positive staining was analyzed (Figure 3G, upper panel) and represented numerically. The expression score is significantly lower in the carcinoma stage as compared with either hyperplasia or adenoma (p < 0.0001, Two-way ANOVA followed with Sidak’s test). There was no significant difference within each lesion type between the two treatment groups for the extent of PR-positivity (Figure 3G, lower panel).

## Discussion

Our results show that when MMTV-PyMT mice are administered UNC10201652, a potent BGUS inhibitor, the incidence of breast tumor (and tumor initiating) lesions ranging from hyperplasia to frank carcinoma is not significantly changed. There is a trend, however small, that the incidence of hyperplastic and adenomas may be decreased in UNC10201652 treated mice; however, as we will discuss this would require a different experimental setup to clarify. To our knowledge this is the first report of use of any BGUS inhibitor in chemoprevention of breast tumors using a genetic model simulating human breast cancer.

There are no mouse transgenic breast cancer models that are truly representative of human ER+ tumors, especially luminal A tumors. The *p53*^+/−^ model developed by Dan Medina at Baylor [45] does closely simulate ER+ breast cancers. Also, mice with mammary gland expression of constitutively active AKT (myrAKT) develop ER+ tumors upon DMBA treatment [46]. To our knowledge, this might be the closest model to human tumors (because PI3K activating mutations are reasonably common in human ER+ cancers, although activating mutations of AKT are less common), however it is quite time-consuming. These models require carcinogen promotion and for purposes of evaluating enzymes that might alter carcinogen metabolism (e.g., microbial BGUS) [47], these models are not tractable for the study of tumor prevention. On the other hand, the MMTV-PyMT tumor model is tractable as it does not involve use of carcinogens to promote tumor formation; relies on a “natural” progression of discrete steps in mammary carcinogenesis; and could potentially be used for the study of male (unpublished observations, Mani Laboratory, Bronx, NY) and female breast cancer development [16].

The results of our study have caveats that raise the need for further experimentation to definitely evaluate the effect of BGUS inhibition in breast tumorigenesis. First, several transgenic and knockout breast cancer models need to be evaluated and their results compared with our MMTV-PyMT mice [48]. It is conceivable that the effect we see if specific to this model. Second, an in-depth evaluation of the level of microbial BGUS inhibition (feces) by UNC10201652 needs to be performed in the MMTV-PyMT mice to determine the extent to which inhibition of all relevant BGUS enzymes occurs at the chosen dose (20 μM/day). It is important to note that this dose was chosen based on prior observations that BGUS inhibition at this dose are effective in vivo across several mouse models [27, 28, 31, 35]. In this context, evaluation of the intestinal and breast microbiota in terms of any major changes in species/class/phyla using 16S or other metagenomic sequencing needs to be carried out. The animal models also need to be evaluated in terms of their metabolomics – in particular, fecal, serum and breast concentration of estrogen and its major metabolites, glucuronides and sulfates. Third, use of systemically bioavailable BGUS inhibitors would be of great use in the study of whether breast nipple fluid BGUS inhibition (resident bacteria in the duct fluid also express BGUS enzymes; changing the nipple aspirate “estrabolome”) also lowers re-absorbed estrogen in the glands and reduces tumor formation locally [5].

Our study shows a minor but potentially interesting trend in the data. We find a trend favoring lower rates of hyperplastic and adenomatous lesions, as well as lower Ki67 and PCNA scores in MMTV-PyMT mice treated with UNC10201652. While this may be a spurious result, these observations may well be correct since our mouse model started chemoprevention at 4 weeks of age. In this model, at 4 weeks, most mice have already initiated tumor development [17]. Indeed, if we know that the highest expression of ER is in the early lesions (hyperplasia, adenoma), then it is likely that bioavailable estrogen could potently drive early lesion formation and/or development [17]. Thus, reducing overall estrogen exposure in breast tissue *early at birth* would be critical in addressing this problem. A re-design of our experiments would be necessary to address protection against the development of hyperplasia/adenoma and several approaches could be considered. One could be the treatment of the mother of the index MMTV-PyMT mouse with BGUS inhibitors that are excreted in mammary milk. The female pups would then be exposed to BGUS inhibitors at birth and through weaning. The other could be attempting to solubilize UNC10201652 in other media such as milk or create tasteless formulations for oral gavage. In this context, perhaps injectable (intravenous, subcutaneous) formulations may also be considered. Thus, overall, formulations for solubility and palatability coupled with early delivery via milk or water is indicated with a study limited to 2-4 weeks.

In the context of using BGUS inhibitors to target chemotherapy toxicity [27], our study provides some valuable groundwork in terms of safety and effects on cancer progression, especially in advanced stages. Our data clearly shows that there is no biologically or statistically significant enhancement of tumor progression or late tumor incidence by UNC10201652. Thus, it is safe to use UNC10201652 in MMTV-PyMT mice in conjunction with other chemotherapeutics in attempts to enhance anti-tumor efficacy while potentially lowering GI toxicity.

---

**Supplementary Figure 1. Genotyping of MMTV-PyMT mice.** The right arrow identifies the index amplicon identifying that mouse as harboring the PyMT antigen gene. M, male; F, female

## Ethical Standards

All the experiments in this manuscript comply with current laws about experimentation in the United States of America. This article does not contain any studies with human participants performed by any of the authors. Appropriate animal studies approvals were obtained from both The Albert Einstein College of Medicine, Bronx, NY and The Army Medical Research and Materiel Command (USAMRMC) Animal Care and Use Review Office (ACURO) all approved ethical/euthanasia guidelines were strictly followed without deviation.

## Conflict of Interest

SM is a voluntary (*non-paid*) scientific advisor to Symberix, INC (https://www.symberix.net/). MRR is the Chief Scientific Officer to Symberix, INC. MRR and Symberix, INC have filed patents on these compounds.

In addition, non-overlapping BGUS inhibitor research has been previously supported by NIH (R01 CA161879 FROM 4/1/2012 To 3/31/2018 to SM and MRR).

## Research Data Policy

All the data presented within this manuscript is freely available for distribution for use in non-commercial purposes. The raw datasets, computation and pathology reports (descriptive) are freely available upon request with the senior authors.

## Acknowledgement

This study was fully supported by The Department of Defense, U.S. Army Medical Research and Materiel Command Congressionally Directed Medical Research Programs, 2016 Breast Cancer Research Program Breakthrough Award - Funding Level 1 (CDMRP Log # BC161093; GRANT12158914; Award# W81XWH-17-1-0023) (PI: SM). We thank the following core services for their assistance in generating the data – The Cancer Center Grant P30CA013330 (PI: David Goldman), The Institute for Animal Studies (Officers: Sunder Shrestha, Jorge Larocca, Linda Jelicks, and Lawrence Herbst) and, The Histology and Comparative Pathology Facility of the Albert Einstein College of Medicine (Director: APB; Technician: Hong Zhang), Bronx, NY 10461.

## References

1. Siegel RL, Miller KD, Jemal A: Cancer statistics, 2015. CA Cancer J Clin 2015, 65(1):5–29.

2. Stierer M, Rosen H, Weber R, Hanak H, Spona J, Tuchler H: Immunohistochemical and biochemical measurement of estrogen and progesterone receptors in primary breast cancer. Correlation of histopathology and prognostic factors. Ann Surg 1993, 218(1):13–21.

3. Carroll JS, Hickey TE, Tarulli GA, Williams M, Tilley WD: Deciphering the divergent roles of progestogens in breast cancer. Nat Rev Cancer 2017, 17(1):54–64.

4. Kwa M, Plottel CS, Blaser MJ, Adams S: The Intestinal Microbiome and Estrogen Receptor-Positive Female Breast Cancer. J Natl Cancer Inst 2016, 108(8).

5. Mani S: Microbiota and Breast Cancer. Prog Mol Biol Transl Sci 2017, 151:217–229.

6. Baker JM, Al-Nakkash L, Herbst-Kralovetz MM: Estrogen-gut microbiome axis: Physiological and clinical implications. Maturitas 2017, 103:45–53.

7. Flores R, Shi J, Fuhrman B, Xu X, Veenstra TD, Gail MH, Gajer P, Ravel J, Goedert JJ: Fecal microbial determinants of fecal and systemic estrogens and estrogen metabolites: a cross-sectional study. J Transl Med 2012, 10:253.

8. Adlercreutz H, Martin F, Pulkkinen M, Dencker H, Rimer U, Sjoberg NO, Tikkanen MJ: Intestinal metabolism of estrogens. J Clin Endocrinol Metab 1976, 43(3):497–505.

9. Heimer GM, Englund DE: Enterohepatic recirculation of oestriol studied in cholecystectomized and non-cholecystectomized menopausal women. Ups J Med Sci 1984, 89(2):107–115.

10. Chen KLA, Liu X, Zhao YC, Hieronymi K, Rossi G, Auvil LS, Welge M, Bushell C, Smith RL, Carlson KE et al: Long-Term Administration of Conjugated Estrogen and Bazedoxifene Decreased Murine Fecal beta-Glucuronidase Activity Without Impacting Overall Microbiome Community. Sci Rep 2018, 8(1):8166.

11. Goldin BR, Adlercreutz H, Gorbach SL, Warram JH, Dwyer JT, Swenson L, Woods MN: Estrogen excretion patterns and plasma levels in vegetarian and omnivorous women. N Engl J Med 1982, 307(25):1542–1547.

12. Chan AA, Bashir M, Rivas MN, Duvall K, Sieling PA, Pieber TR, Vaishampayan PA, Love SM, Lee DJ: Characterization of the microbiome of nipple aspirate fluid of breast cancer survivors. Sci Rep 2016, 6:28061.

13. Cohen LA, Zhao Z, Zang EA, Wynn TT, Simi B, Rivenson A: Wheat bran and psyllium diets: effects on N-methylnitrosourea-induced mammary tumorigenesis in F344 rats. J Natl Cancer Inst 1996, 88(13):899–907.

14. Zhang J, Lacroix C, Wortmann E, Ruscheweyh HJ, Sunagawa S, Sturla SJ, Schwab C: Gut microbial beta-glucuronidase and glycerol/diol dehydratase activity contribute to dietary heterocyclic amine biotransformation. BMC Microbiol 2019, 19(1):99.

15. Humblot C, Murkovic M, Rigottier-Gois L, Bensaada M, Bouclet A, Andrieux C, Anba J, Rabot S: Beta-glucuronidase in human intestinal microbiota is necessary for the colonic genotoxicity of the food-borne carcinogen 2-amino-3-methylimidazo[4,5-f]quinoline in rats. Carcinogenesis 2007, 28(11):2419–2425.

16. Fluck MM, Schaffhausen BS: Lessons in signaling and tumorigenesis from polyomavirus middle T antigen. Microbiol Mol Biol Rev 2009, 73(3):542–563, Table of Contents.

17. Lin EY, Jones JG, Li P, Zhu L, Whitney KD, Muller WJ, Pollard JW: Progression to malignancy in the polyoma middle T oncoprotein mouse breast cancer model provides a reliable model for human diseases. Am J Pathol 2003, 163(5):2113–2126.

18. Cowen S, McLaughlin SL, Hobbs G, Coad J, Martin KH, Olfert IM, Vona-Davis L: High-Fat, High-Calorie Diet Enhances Mammary Carcinogenesis and Local Inflammation in MMTV-PyMT Mouse Model of Breast Cancer. Cancers (Basel) 2015, 7(3):1125–1142.

19. Rossdeutscher L, Li J, Luco AL, Fadhil I, Ochietti B, Camirand A, Huang DC, Reinhardt TA, Muller W, Kremer R: Chemoprevention activity of 25-hydroxyvitamin D in the MMTV-PyMT mouse model of breast cancer. Cancer Prev Res (Phila) 2015, 8(2):120–128.

20. Gluschnaider U, Hertz R, Ohayon S, Smeir E, Smets M, Pikarsky E, Bar-Tana J: Long-chain fatty acid analogues suppress breast tumorigenesis and progression. Cancer Res 2014, 74(23):6991–7002.

21. Gil EY, Jo UH, Lee HJ, Kang J, Seo JH, Lee ES, Kim YH, Kim I, Phan-Lai V, Disis ML et al: Vaccination with ErbB-2 peptides prevents cancer stem cell expansion and suppresses the development of spontaneous tumors in MMTV-PyMT transgenic mice. Breast Cancer Res Treat 2014, 147(1):69–80.

22. Wang C, Schwab LP, Fan M, Seagroves TN, Buolamwini JK: Chemoprevention activity of dipyridamole in the MMTV-PyMT transgenic mouse model of breast cancer. Cancer Prev Res (Phila) 2013, 6(5):437–447.

23. Ma J, Lanza DG, Guest I, Uk-Lim C, Glinskii A, Glinsky G, Sell S: Characterization of mammary cancer stem cells in the MMTV-PyMT mouse model. Tumour Biol 2012, 33(6):1983–1996.

24. Christenson JL, Butterfield KT, Spoelstra NS, Norris JD, Josan JS, Pollock JA, McDonnell DP, Katzenellenbogen BS, Katzenellenbogen JA, Richer JK: MMTV-PyMT and Derived Met-1 Mouse Mammary Tumor Cells as Models for Studying the Role of the Androgen Receptor in Triple-Negative Breast Cancer Progression. Horm Cancer 2017, 8(2):69–77.

25. Park J, Kusminski CM, Chua SC, Scherer PE: Leptin receptor signaling supports cancer cell metabolism through suppression of mitochondrial respiration in vivo. Am J Pathol 2010, 177(6):3133–3144.

26. Wallace BD, Redinbo MR: The human microbiome is a source of therapeutic drug targets. Curr Opin Chem Biol 2013, 17(3):379–384.

27. Wallace BD, Wang H, Lane KT, Scott JE, Orans J, Koo JS, Venkatesh M, Jobin C, Yeh LA, Mani S et al: Alleviating cancer drug toxicity by inhibiting a bacterial enzyme. Science 2010, 330(6005):831–835.

28. Roberts AB, Wallace BD, Venkatesh MK, Mani S, Redinbo MR: Molecular insights into microbial beta-glucuronidase inhibition to abrogate CPT-11 toxicity. Mol Pharmacol 2013, 84(2):208–217.

29. Pellock SJ, Redinbo MR: Glucuronides in the gut: Sugar-driven symbioses between microbe and host. J Biol Chem 2017, 292(21):8569–8576.

30. Wallace BD, Roberts AB, Pollet RM, Ingle JD, Biernat KA, Pellock SJ, Venkatesh MK, Guthrie L, O’Neal SK, Robinson SJ et al: Structure and Inhibition of Microbiome beta-Glucuronidases Essential to the Alleviation of Cancer Drug Toxicity. Chem Biol 2015, 22(9):1238–1249.

31. LoGuidice A, Wallace BD, Bendel L, Redinbo MR, Boelsterli UA: Pharmacologic targeting of bacterial beta-glucuronidase alleviates nonsteroidal anti-inflammatory drug-induced enteropathy in mice. J Pharmacol Exp Ther 2012, 341(2):447–454.

32. Boelsterli UA, Redinbo MR, Saitta KS: Multiple NSAID-induced hits injure the small intestine: underlying mechanisms and novel strategies. Toxicol Sci 2013, 131(2):654–667.

33. Saitta KS, Zhang C, Lee KK, Fujimoto K, Redinbo MR, Boelsterli UA: Bacterial beta-glucuronidase inhibition protects mice against enteropathy induced by indomethacin, ketoprofen or diclofenac: mode of action and pharmacokinetics. Xenobiotica 2014, 44(1):28–35.

34. Mani S, Boelsterli UA, Redinbo MR: Understanding and modulating mammalian-microbial communication for improved human health. Annu Rev Pharmacol Toxicol 2014, 54:559–580.

35. Yauw STK, Arron M, Lomme R, van den Broek P, Greupink R, Bhatt AP, Redinbo MR, van Goor H: Microbial Glucuronidase Inhibition Reduces Severity of Diclofenac-Induced Anastomotic Leak in Rats. Surg Infect (Larchmt) 2018, 19(4):417–423.

36. Pellock SJ, Creekmore BC, Walton WG, Mehta N, Biernat KA, Cesmat AP, Ariyarathna Y, Dunn ZD, Li B, Jin J et al: Gut Microbial beta-Glucuronidase Inhibition via Catalytic Cycle Interception. ACS Cent Sci 2018, 4(7):868–879.

37. Biernat KA, Pellock SJ, Bhatt AP, Bivins MM, Walton WG, Tran BNT, Wei L, Snider MC, Cesmat AP, Tripathy A et al: Structure, function, and inhibition of drug reactivating human gut microbial beta-glucuronidases. Sci Rep 2019, 9(1):825.

38. Pellock SJ, Walton WG, Biernat KA, Torres-Rivera D, Creekmore BC, Xu Y, Liu J, Tripathy A, Stewart LJ, Redinbo MR: Three structurally and functionally distinct beta-glucuronidases from the human gut microbe Bacteroides uniformis. J Biol Chem 2018, 293(48):18559–18573.

39. Venkatesh M, Mukherjee S, Wang H, Li H, Sun K, Benechet AP, Qiu Z, Maher L, Redinbo MR, Phillips RS et al: Symbiotic bacterial metabolites regulate gastrointestinal barrier function via the xenobiotic sensor PXR and Toll-like receptor 4. Immunity 2014, 41(2):296–310.

40. Guy CT, Cardiff RD, Muller WJ: Induction of mammary tumors by expression of polyomavirus middle T oncogene: a transgenic mouse model for metastatic disease. Molecular and Cellular Biology 1992, 12(3):954.

41. Dowsett M, Nielsen TO, A’Hern R, Bartlett J, Coombes RC, Cuzick J, Ellis M, Henry NL, Hugh JC, Lively T et al: Assessment of Ki67 in breast cancer: recommendations from the International Ki67 in Breast Cancer working group. J Natl Cancer Inst 2011, 103(22):1656–1664.

42. Zhang D, Leal AS, Carapellucci S, Zydeck K, Sporn MB, Liby KT: Chemoprevention of Preclinical Breast and Lung Cancer with the Bromodomain Inhibitor I-BET 762. Cancer Prev Res (Phila) 2018, 11(3):143–156.

43. Faul F, Erdfelder E, Lang AG, Buchner A: G*Power 3: a flexible statistical power analysis program for the social, behavioral, and biomedical sciences. Behav Res Methods 2007, 39(2):175–191.

44. Duivenvoorden HM, Spurling A, O’Toole SA, Parker BS: Discriminating the earliest stages of mammary carcinoma using myoepithelial and proliferative markers. PLoS One 2018, 13(7):e0201370.

45. Jerry DJ, Kittrell FS, Kuperwasser C, Laucirica R, Dickinson ES, Bonilla PJ, Butel JS, Medina D: A mammary-specific model demonstrates the role of the p53 tumor suppressor gene in tumor development. Oncogene 2000, 19(8):1052–1058.

46. Blanco-Aparicio C, Pérez-Gallego L, Pequeño B, Leal JFM, Renner O, Carnero A: Mice expressing myrAKT1 in the mammary gland develop carcinogen-induced ER-positive mammary tumors that mimic human breast cancer. Carcinogenesis 2007, 28(3):584–594.

47. Vater ST, Baldwin DM, Warshawsky D: Hepatic metabolism of 7,12-dimethylbenz(a)anthracene in male, female, and ovariectomized Sprague-Dawley rats. Cancer Res 1991, 51(2):492–498.

48. Swiatnicki MR, Andrechek ER: How to Choose a Mouse Model of Breast Cancer, a Genomic Perspective. J Mammary Gland Biol Neoplasia 2019.

