## Supplementary figures and images for "Inhibition of Microbial Beta-Glucuronidase Does Not Prevent Breast Carcinogenesis in the Polyoma Middle T Mouse"

### Supplemental Figure 1

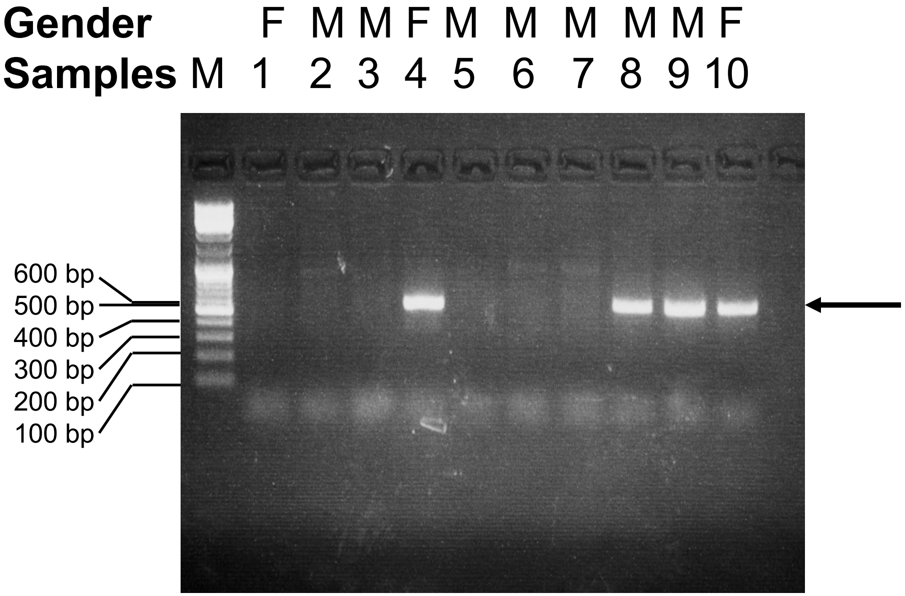
